# Prognostic Analysis of Histopathological Images Using Pre-Trained Convolutional Neural Networks

**DOI:** 10.1101/620773

**Authors:** Liangqun Lu, Bernie Daigle

**Affiliations:** Department of Biological Sciences, University of Memphis; Department of Computer Science, University of Memphis

**Keywords:** Histopathological Images, Convolutional Neural Networks, Hepatocellular Carcinoma, Feature Extraction

## Abstract

**Background:** Histopathological images contain rich phenotypic descriptions of the molecular processes underlying disease progression. Convolutional neural networks (CNNs), a state-of-the-art image analysis technique in computer vision, automatically learns representative features from such images which can be useful for disease diagnosis, prognosis, and subtyping. Despite hepatocellular carcinoma (HCC) being the sixth most common type of primary liver malignancy with a high mortality rate, little previous work has made use of CNN models to delineate the importance of histopathological images in diagnosis and clinical survival of HCC.

**Results:** We applied three pre-trained CNN models – VGG 16, Inception V3, and ResNet 50 – to extract features from HCC histopathological images. The visualization and classification showed clear separation between cancer and normal samples using image features. In a univariate Cox regression analysis, 21.4% and 16% of image features on average were significantly associated with overall survival and disease-free survival, respectively. We also observed significant correlations between these features and integrated biological pathways derived from gene expression and copy number variation. Using an elastic net regularized CoxPH model of overall survival, we obtained a concordance index (C-index) of 0.789 and a significant log-rank test (p = 7.6E-18) after applying Inception image features. We also performed unsupervised classification to identify HCC subgroups from image features. The optimal two subgroups discovered using Inception image features were significantly associated with both overall (C-index = 0.628 and p = 7.39E-07) and disease-free survival (C-index =0.558 and p = 0.012). Our results suggest the feasibility of feature extraction using pre-trained models, as well as the utility of the resulting features to build an accurate prognosis model of HCC and highlight significant correlations with clinical survival and biological pathways.

**Conclusions:** The image features extracted from HCC histopathological images using the pre-trained CNN models VGG 16, Inception V3 and ResNet 50 can accurately distinguish normal and cancer samples. Furthermore, these image features are significantly correlated with relevant biological outcomes.

## Background

Histopathological images contain rich phenotypic descriptions of the molecular processes underlying disease progression and have been used for diagnosis, prognosis, and subtype discovery [1]. These images contain visual features including nuclear atypia, mitotic activity, cellular density, tissue architecture and higher-order patterns, which are examined by pathologists to diagnose and grade lesions. The recentaccumulated scanned whole slide images (WSI) have boosted the wide application of machine learning algorithms to extract useful information and assist tasks in the areas of lesion detection, classification, segmentation, and image reconstruction [2]. Deep learning is a broad family of machine learning methods based on deep neural network representations, which have been widely applied in recent computer vision and natural language processing tasks [3]. Convolutional neural networks (CNNs), a state-of-the-art image analysis technique in computer vision, automatically learns representative features from images and has been dominant since its astonishing results at the ImageNet Large Scale Visual Recognition Competition (ILSVRC) in 2012 [4]. In various studies, CNNs have shown good performance when applied to medical images, including those from radiology [5] [6] [7]. Additional studies in the areas of diabetic retinopathy screening [8], skin lesion classification [9], and lymph node metastasis detection [10] have demonstrated expert-level performance by CNNs. Compared with traditional machine learning techniques, CNNs have witnessed significant advances in areas of image registration and localization, detection of anatomical and cellular structures, tissue segmentation, and computer-aided disease prognosis and diagnosis [11].

Primary liver cancer is the 6th most common type of liver malignancy with a high mortality and morbidity rate. Hepatocellular carcinoma (HCC) is the representative type resulting from the malignant transformation of hepatocytes in a cirrhotic, non-fibrotic, or minimal fibrotic liver [12]. With the development of high-throughput technologies, a number of “omics” research studies have helped detail the mechanism of molecular pathogenesis, which has significantly contributed to our understanding of cancer genomics, diagnostics, prognostics, and therapy in an unprecedented way [13] [14] [15] [16]. The most frequent mutations and chromosome alterations leading to HCC were identified in the TERT promoter, as well as the CTNNB1, TP53, AXIN1, ARID1A, NFE2L2, ARID2 and RPS6KA3 genes [16]. The biological pathways Wnt/*β*-catenin signaling, oxidative stress metabolism, and Ras/mitogen-activated protein kinase (MAPK) were reported to be involved in liver carcinogenesis [13]. In addition, frequent TP53-inactivating mutations, higher expression of stemness markers (KRT19, EPCAM) and the tumor marker BIRC5, and activated Wnt and Akt signaling pathways were reported to associate with stratification of HCC samples [16]. Furthermore, histological subtypes of HCC have been shown to be related to particular gene mutations and molecular tumour classification [17]. Two recent studies have demonstrated strong connections between molecular changes and disease phenotypes. In a meta-analysis of 1494 HCC samples, consensus driver genes were identified that showed strong impacts on cancer phenotypes [18]. In addition, a deep learning-based multi-omics integration study produced a model capable of robust survival prediction [19]. These and other recent findings may help to translate our knowledge of HCC biology into clinical practice [17].

At the pathological level, HCC exhibits as a morphologically heterogeneous tumour. Although HCC neoplastic cells most often grow in cords of variable thickness lined by endothelial cells mimicking the trabeculae and sinusoids of normal liver, other architectural patterns are frequently observed and numerous cytological variants recognized, including clear, pleomorphic, or bile producing cells. Though histopathologic criteria for diagnosing classical, progressed HCC are well established and known, it is challenging to detect increasingly small lesions in core needle biopsies during routine screening diagnosis programs. These lesions can be far more difficult to distinguish from one another than progressed HCC, which is usually diagnosed in a clear cut manner using hematoxylin and eosin staining [20] [21]. Although prognostication increasingly relies on genomic biomarkers that measure genetic alterations, gene expression changes, and epigenetic modifications, histology remains an important tool in predicting the future course of a patient’s disease. Previous studies [22] [23] indicate the complementary information between histopathological and genomic data. Quantitative analysis of these images and their integration with genomics data require innovations in integrative genomics and call for techniques from bioimage informatics, genomics, and bioinformatics.

In this study, we applied pre-trained CNN models on HCC histopathological images to extract image features and characterize the relationships between images, clinical survival and biological pathways in order to leverage both modalities to improve patient outcomes eventually. We downloaded Hematoxylin and eosin (HE) - stained whole-slide images from HCC subjects (421 tumor samples and 105 normal tissue adjacent to tumor samples) from the National Cancer Institute Genomic Data Commons Data Portal. We then applied three pre-trained CNN models–VGG 16, Inception V3, and ResNet–to extract features after image normalization. We performed classification between cancer and normal samples using the image features. We also constructed models associating image features with clinical survival. Finally, we calculated correlations between image features and integrated biological pathways. To the best of our knowledge, this is the first study to extract HCC image features using pre-trained CNN models, and our results indicate both the feasibility of CNN model application to histopathological images as well as the relevance of such images to disease survival and biological pathways.

## Results

In this study, we made used of pre-trained CNN models VGG 16, Inception V3 and ResNet 50 to extract features from HCC histopathological whole slide images. Using the image features, we performed survival analysis and subgroup discovery. We also performed correlation analysis between image features and integrated biological pathways. The workflow of analysis steps can be seen in Figure 1.

**Figure 1.**
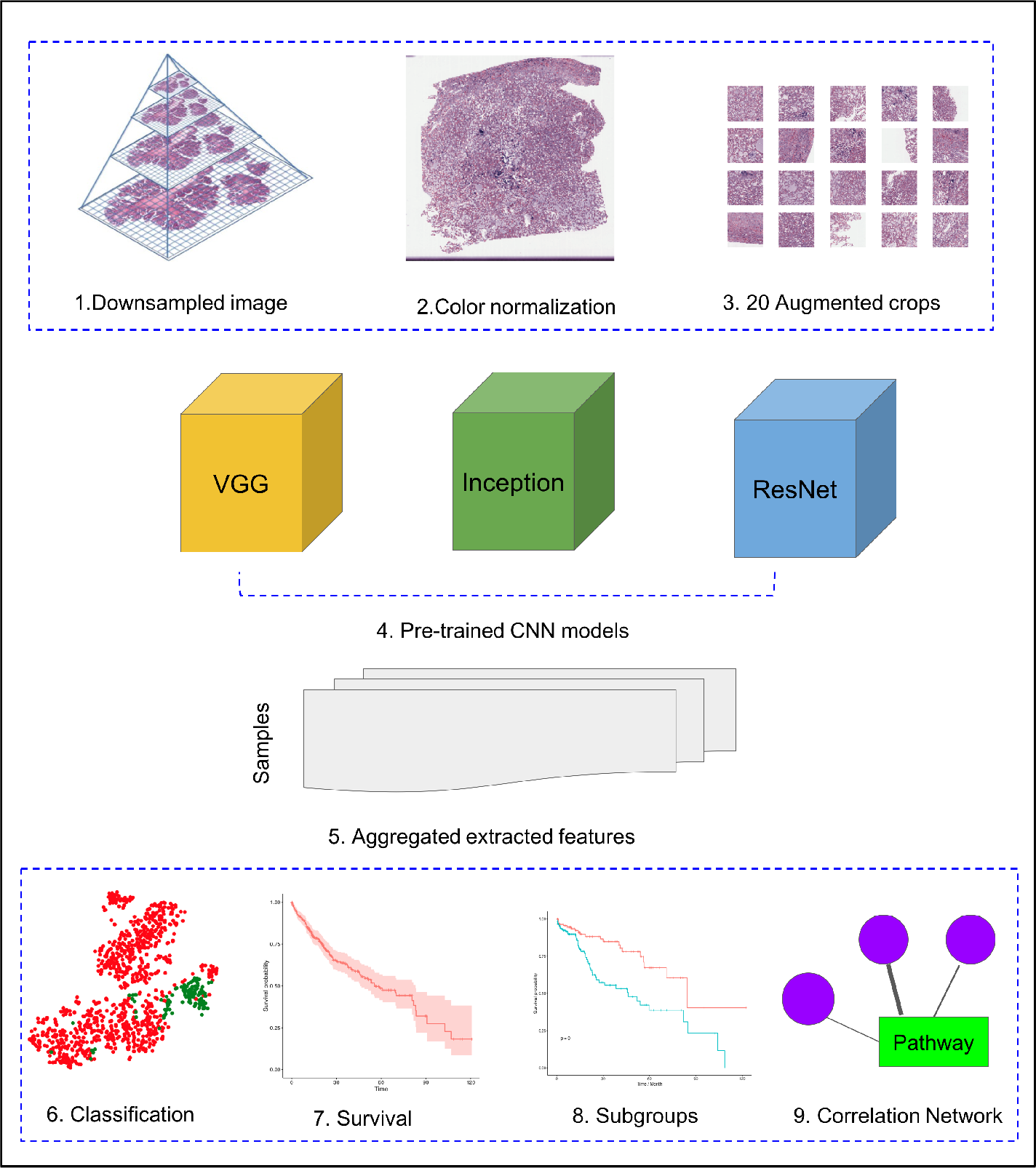
HCC image analysis flow. 1) For whole slide .svs files, downsampled images were generated, 2) color normalization was performed, 3) 50 augmented images were made for each original image and 20 crops were selected at random from each augmented image, 4) three CNN models, VGG 16, Inception 3 and ResNet 50 were applied to extract features from each crop, 5) features from all crops were aggregated and 50 sets of image features were obtained from each CNN model, 6-9) image features were used for the analysis.

### Image feature extraction and survival analysis

Histopathology assessment is mandatory in HCC diagnosis [24] and the characteristics tumor number, size, cell differentiation and grade, and presence of satellite nodules were reported to be prognostic biomarkers [25]. CNNs have the advantage of automatic feature representation using convolution layers, pooling layers, and fully connected layers. In order to examine the image features, we first downloaded HCC histopathological images from the National Cancer Institute Genomic Data Commons Data Portal, in which multiple molecular datasets and clinical information are also available for the same cohort. There were a total of 966 .svs image files from 421 cancer tissues and 105 tumor-adjacent normal tissues, of which 964 had enough information for the following analysis. For all image files, we downsampled and obtained image files with median 5601 × 2249.5 pixels. We performed color normalization and generated 50 images using color augmentation in order to improve sample variety. We randomly selected 20 crops of sizes 512 × 512 pixels or 256 × 256 pixels from each augmented image. The 20 512 × 512 crops represent 41.6% of the input image pixels on average, while the 20 256 × 256 crops represent 10.4% on average. The deep CNN models VGG 16, Inception V3 and ResNet 50 contain millions of parameters, and extensive training of these models has led to state-of-the-art performance in image recognition and classification [26]. We extracted features from these models in an unsupervised way to avoid the challenges of CNN model training from scratch. For the Inception and ResNet models, the second-to-last layers were selected as features after the exclusion of the last fully-connected layers of the networks. For the VGG model, the last four layers were concatenated for features. For each model, we combined features from the 20 random crops into a single set of features to represent each image.

On each of the 50 augmented images, we obtained 1408, 2408, and 2408 features from the VGG 16, Inception V3, and ResNet 50 models, respectively. To aggregate these features across the augmented images, we computed median values for each feature. To facilitate visualization of cancer and normal samples, we first used PCA to reduce the feature dimensionality followed by t-SNE applied to the first 10 principal components. We also performed supervised classification of the samples using a linear Support Vector Machine applied to each set of image features. Figure 2 shows these results using features derived from 256 × 256 crop sizes, with classification performance displayed as receiver operating characteristic (ROC) and two-class precision-recall curves. The average AUC achieved by all three models is between 0.99 and 1, illustrating the clear separation achieved between tumor and normal samples using the extracted image features. The AUCs achieved for features derived from 512 × 512 crop sizes were similarly very close to 1. To compare this performance with that of an alternate method, we also applied PCA (randomized SVD) and SVD (full SVD) on the downsampled images without augmentation. Specifically, we extracted the first 100 principal components (PCA) or singular vectors (SVD) as features and performed supervised classification. Figure S1 shows that performance using PCA- and SVD-derived features are very poor. Finally, we performed classification on features derived without using image augmentation. Here, performance is only slightly worse, with AUCs ranging between 0.98 and 0.99 (Figure S2). Overall, our results show that use of CNN-derived image features is extremely effective for distinguishing HCC tumor from normal samples, which suggests that pre-trained CNN models capture the most relevant characteristics from HCC histopathological images.

**Figure 2.**
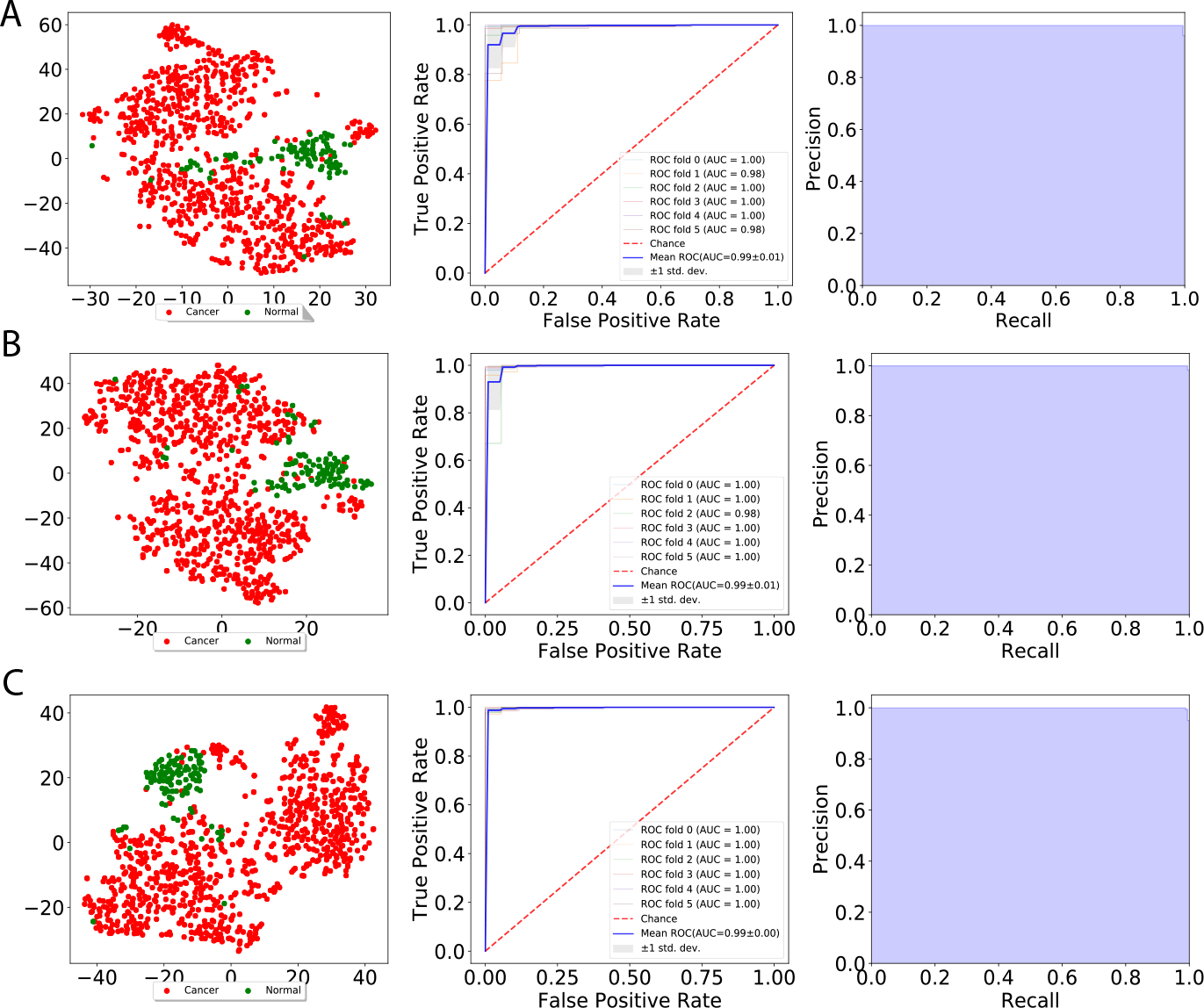
Visualization of extracted image features and classification between cancer and normal samples. The left side indicates t-SNE visualization, the middle indicates ROC curves from linear SVM and the right side indicates Recall and Precision curves measured using A) VGG image features, B) Inception features, C) ResNet features.

In order to illustrate whether the CNN-derived image features are associated with clinical survival, we performed CoxPH regression analyses to predict survival. We obtained clinical information for each sample from the cBioPortal for Cancer Genomics, as described in the Methods section. For samples with more than one image, we computed median feature values across the images. We considered both overall survival (OS) and disease free survival (DFS). In OS, the censored time is the last day of contact, while the censored time for DFS is determined by the nonexistence of new tumors. For each image feature, we applied CoxPH regression models for both OS and DFS and selected significantly associated features (p-value ≤ 0.05) based on a Score (logrank) test. Table 1 shows the number of significant features for each model and survival type. Each model had a slightly different number of features, with more significant features associated with OS than DFS.

**Table 1.** Significant image feature number from univariate CoxPH regression models.

| Model | Feature Number | Crop Size | Significant Features of OS | Significant Features of DFS |
| --- | --- | --- | --- | --- |
| VGG | 1408 | 256 | 272 (19.32%) | 219 (15.55%) |
| Inception | 2048 | 256 | 574 (28.03%) | 294 (14.36%) |
| ResNet | 2048 | 256 | 522 (25.49%) | 385 (18.80%) |
| VGG | 1408 | 512 | 300 (21.31%) | 201 (14.28%) |
| Inception | 2048 | 512 | 356 (17.38%) | 290 (14.16%) |
| ResNet | 2048 | 512 | 347 (16.94%) | 390 (19.04%) |
|  |  |  | average 21.41% | average 16.03% |

Next, we performed multivariate CoxPH regression analyses for each survival type on all image features from each model. We employed elastic net regularization using equal parts of lasso and ridge regularization during model training. Optimal hyper-parameters were selected using 10-fold cross-validation and then used for model prediction. Overall, we identified three multivariate survival models with log-rank p values 1.2e23 (VGG), 7.6e18 (Inception), and 1.2e12 (ResNet), whose survival curves are shown in 3 based on 256 × 256 crop sizes. We also computed the C-index, with the Inception-derived model reaching the highest value of 0.789. Overall, our results show that CNN-derived image features are significantly associated with clinical survival and can be used to build accurate survival models.

### Subgroup discovery from image features

In order to investigate whether our CNN-derived image features relate to HCC prognosis, we next used these features to discover subgroups within tumor samples. We considered all image features which were significantly associated with both OS and DFS. Using these features, we clustered the tumor samples using K-means (K = 2-12) and used both silhouette and Davies-Bouldin values to choose the optimal number of subgroups. As shown in Figure 4, 2 subgroups were determined to be optimal for all three models. We visualized these subgroups using t-SNE to reduce dimensionality.

**Figure 3.**
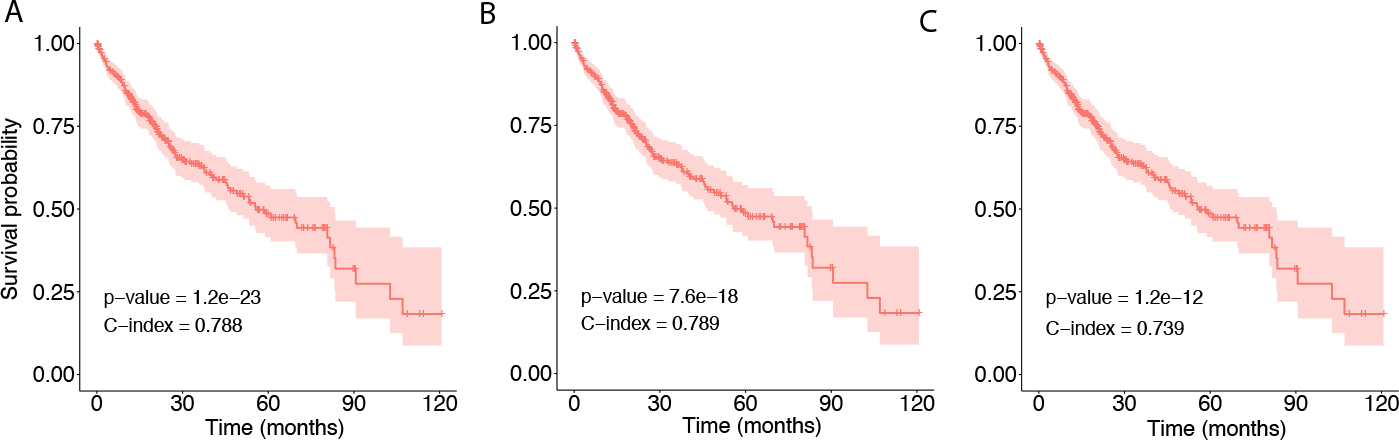
Overall Survival probability from multivariate CoxPH model using 256 × 256 pixel crop size. A) Using VGG image features, B) using Inception image features, C) using ResNet image features.

**Figure 4.**
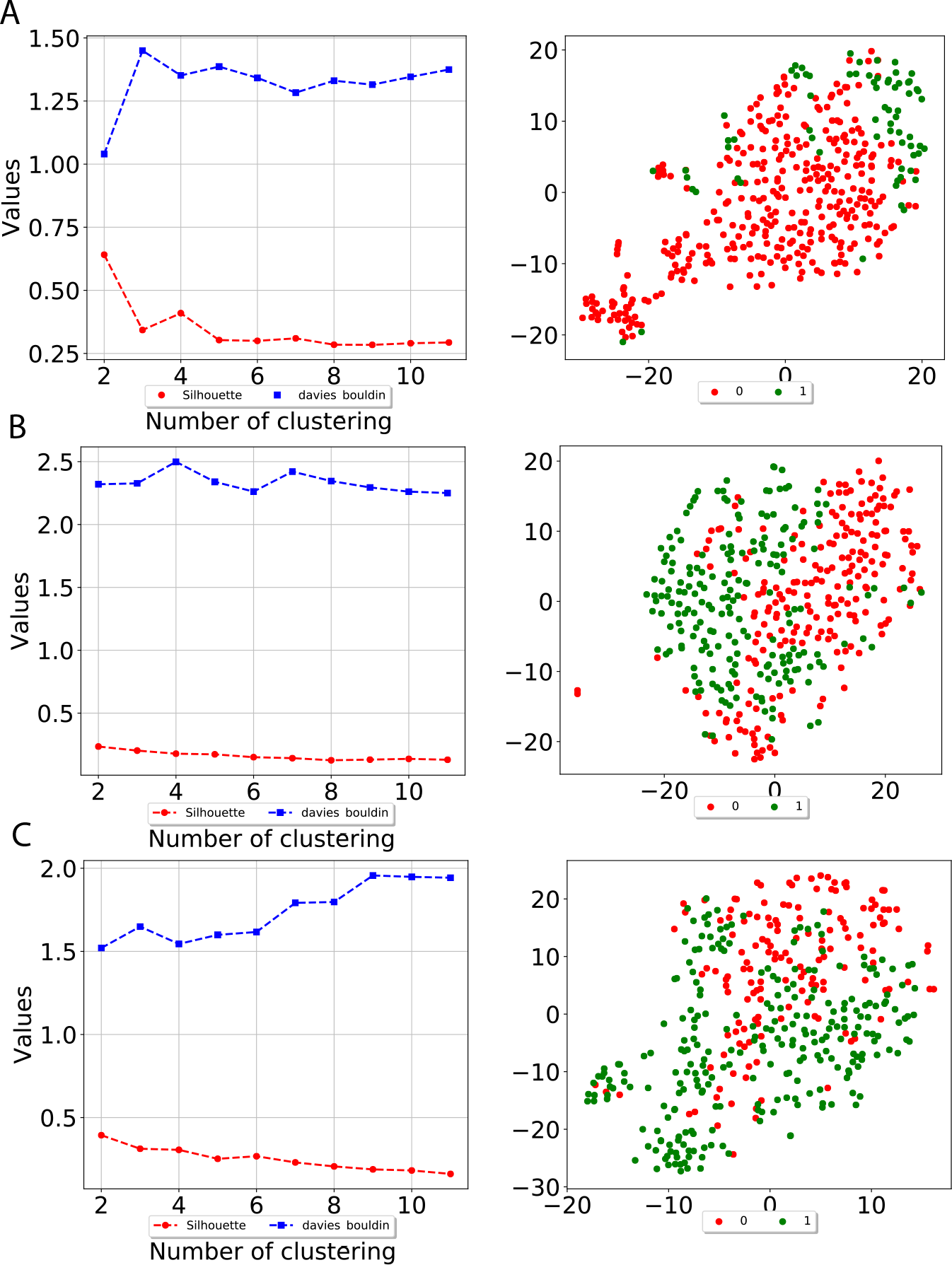
Subgroup discovery from image features using 256 × 256 pixel crop size. The left side displays two different metrics for selecting the optimal number of clusters, and the right side indicates the t-SNE visualization of best clusters. A) using VGG image features, B) using Inception image features, C) using ResNet image features.

We next examined survival differences between the subgroups. For each model and survival type, we applied separate CoxPH models for each subgroup. Figure 5 shows our results, where the subgroups discovered using the Inception and ResNet models both show a significant difference in both OS and DFS using log-rank test. The group 0 is consistently worse in both OS and DFS from Inception model while the group 1 is worse than group 0 at ResNet model. For the VGG model, we only detected a significant difference for DFS. Table 2 shows the subgroup overlap between the three models.

**Figure 5.**
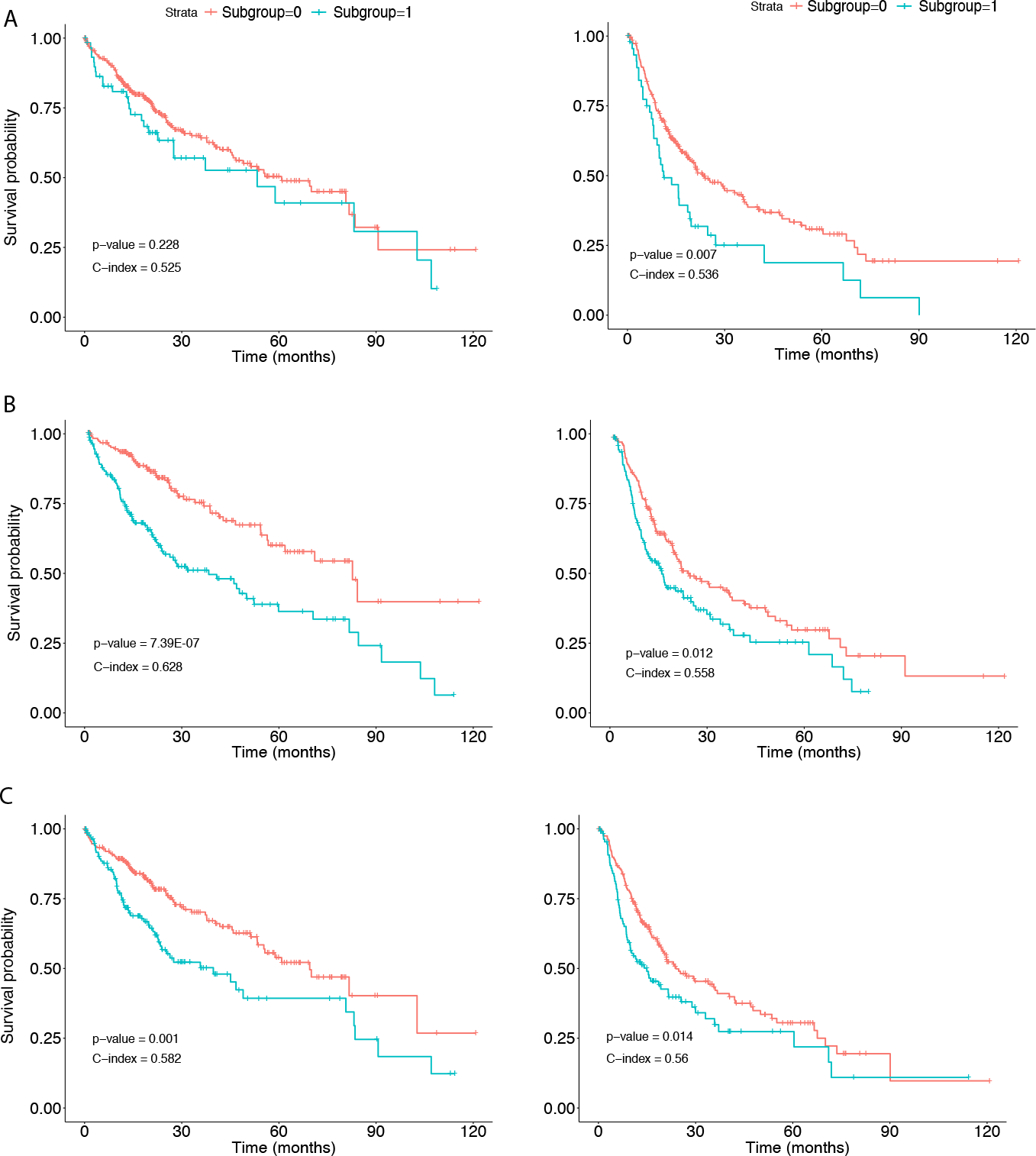
Survival analysis from discovered subgroups. The left side corresponds to the CoxPH model applied to OS, the right side corresponds to DFS. The two groups are indicated in red and green, A) using VGG image features, B) using Inception image features, C) using ResNet image features.

**Table 2.** Overlaps of subgroup frequency counts between three models.

|  |  | Inception | 0 | 1 |
| --- | --- | --- | --- | --- |
| VGG | ResNet |  |  |  |
| 0 | 0 |  | 176 | 48 |
|  | 1 |  | 18 | 109 |
| 1 | 0 |  | 20 | 16 |
|  | 1 |  | 4 | 30 |

### Correlation between image features and biological pathways

Previous studies examined the molecular mechanisms of HCC [13] [14] [15] [16]. To relate our CNN-derived image features to such mechanisms, we identified correlations between features and a collection of molecular pathways. Specifically, we first obtained integrated pathway levels (IPLs) using the Firehose Genome Browser, which provides analysis-ready files inferred from both gene expression and DNA copy number variation using the PARADIGM algorithm [27]. IPLs indicate the predicted activities of biological concepts using both copy number and gene expression data, as described in the Methods section. The total 7203 entities of the IPL matrix are from 3656 concepts in 135 merged pathways, denoted such as `19_EPHB3` – the concept (gene) EPHB3 participating in EPHB forward signaling with pathway index 19. We then computed Pearson correlation coefficients between these IPLs and each feature significantly associated with both OS and DFS. We selected significantly correlated IPL-feature pairs based on Benjamini Hochberg (BH) [28]-adjusted p-value ≤ 0.05. With 256 × 256 crop sizes, 90 (out of 97), 199 (out of 203) and 192 (out of 203) image features were significantly correlated with biological pathways from the VGG, Inception, and ResNet models, respectively. On average, 90.2% of the image features showed a significant correlation, with Pearson correlation coefficients ranging between −0.536 and 0.385.

Finally, we performed differential expression analysis to identify IPL differences between each pair of subgroups. For each model, we selected pathways with BH-adjusted p-values 0.05. Surprisingly, we found no significant pathways at this threshold for all three models and both crop sizes. After relaxing the p-value threshold to 0.1, we detected five significant entities from 2 pathways: EPHB forward signaling (EPHB3, ROCK1, Ephrin B1/EPHB3) and 66: Glucocorticoid receptor regulatory network (IL8, ICAM1). Figure 6 shows a network visualization of these pathways with significantly-correlated image features. Overall, 31 out of 49 image features with significant correlations were found using the Inception model.

**Figure 6.**
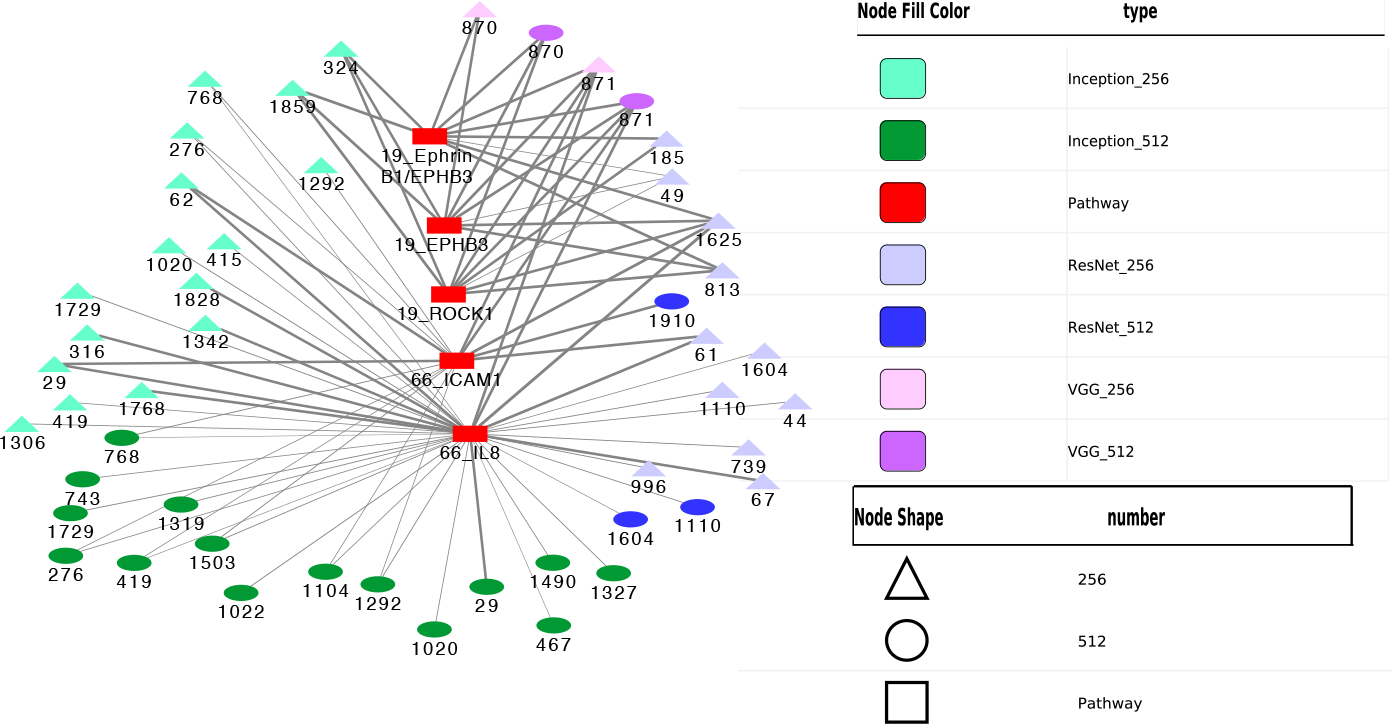
Correlation network between image features and example pathways. Colors of nodes indicate CNN models VGG, Inception and ResNet, as well as pathways. The labeled names of image features consist of the model name, crop size and feature order number. The thickness of each edge corresponds to the magnitude of correlation coefficients ranging between −0.536 and 0.385 that were statistically significant with the range.

## Discussion

In this study, we applied the pre-trained CNN models VGG 16, Inception V3, and ResNet 50 to extract features from HCC histopathological whole slide images. Using these image features, we obtained clear separation between tumor and normal samples in both t-SNE visualization and supervised classification. When considering associations with overall (OS) and disease-free survival (DFS), averages of 21.4% and 16% of image features, respectively, were significant based on univariate CoxPH regression analyses. These image features demonstrated significance in multivariate CoxPH regression OS model. We utilized these image features to discover HCC subgroups, and the resulting subgroups showed a significant difference in survival. Furthermore, 90.2% of the image features were significantly correlated with measures of integrated pathway levels on average. The five significant entities from 2 pathways – EPHB forward signaling and Glucocorticoid receptor regulatory network – implied a potential role for these entities in determining the prognosis of HCC. EPHB forward signaling induces cell repulsion and controls actin cell adhesion and migration [29]. It has been reported that EPHB receptors and ephrin ligands are involved in carcinogenesis and cancer progression [30] and EPHB3 receptor inhibits Wnt signaling pathway [31], which were reported to be useful for HCC stratification [16]. Previous studies have reported that the glucocorticoid receptor (GR) binds promoters, interacts with other transcription factors [32], and the impairment of GR signaling causes hepatocellular carcinoma [33] in mice. IL 8, also known as Interleukin-8, a proinflammatory CXC chemokine, was reported to promote malignant cancer progression [34]. ICAM-1, also known as Intercellular cell adhesion molecule-1, has functions in immune and inflammatory responses and was reported to play a role in liver metastasis [35].

CNNs have shown impressive improvements in deep feature representation learning from medical images in applications such as image classification, image segmentation, and computer-aided disease diagnosis/prognosis [6][36]. As one class of deep learning models, CNNs require massive amounts of data, which can be a challenge for biomedical image analysis studies. Furthermore, deep feature learning depends on the size and degree of annotation of images, which are often not standardized across different datasets. One possible solution for image datasets with a small sample size is transfer learning in which pre-trained CNN models from large-scale natural image datasets were applied to solve biomedical image tasks. In a previous study of CNN models applied to thoraco-abdominal lymph node detection and interstitial lung disease classification, transfer-learning from the large scale annotated image datasets (ImageNet) was consistently beneficial in both tasks [37].

In the application of histopathological image analysis, the large image size and different levels of resolutions from the whole slide images (WSIs) pose challenges to sufficient distinction information selection, and the computational power and training time [2]. In order to avoid the information loss, WSIs are often divided into small patches (ex: 256 × 256 pixels) and each patch is analyzed individually as Region of interest (ROI). These ROIs can also be labeled using active learning [38] or by professional trained pathologists [39]. Then the integrated patch-level decision or object-level decision from averaging regions of patches representing WSIs are studied for the specific tasks [2]. In out work, we applied CNN models with pre-trained weights from the large scale ImageNet dataset. For randomly selected 20 patches of 256 × 256 and 512 × 512 pixels from WSI, we extracted features from the last layers of CNN models to represent each image for visualization and classification. To robustly deal with the color variation and image artifact issues, we conducted color normalization and augmentation in order to remove color variation effects before applying CNN models. Color normalization adjusts pixel-level image values [40], and color augmentation generates more data by altering hue and contrast in the raw images [41]. We achieved very good classification performance, with AUCs between 0.99 and 1 for distinguishing normal and tumor samples. Comparing this performance to previous work, we note that in one study of histopathology images [42], classification performance reached 81.14% accuracy using the extracted features from a pre-trained VGG 19 (similar to VGG 16) network. In a similar study of histopathological images of breast cancer [43], classification performance on 400 HE-stained images of 2048 1536 pixels each reached an AUC of 0.963 for distinguishing between non-carcinomas vs. carcinomas samples. In our study, we have larger histopathological images with median 5601 × 2249.5 pixels indicating more sample information which may explain the better performance.

Stratification of patients is an important step to better understand disease mechanisms and ultimately enable personalized medicine. Previous studies of HCC have suggested molecular-level subgroups [44] [45] [19]. In the latter study, the authors applied deep learning to integrate 3 omic datasets from 360 HCC patients (the same cohort used in our study), discovering two subgroups with survival differences.

In our work, we identified subgroups using all three CNN models, with the subgroups from both Inception (C-index = 0.628; P value = 7.39E-07) and ResNet (C-index = 0.582; P value = 0.001) models showing significant differences in OS. We note that this significance of the Inception model is lower than that achieved using subgroups identified using multiple omic data integration (C-index = 0.68 and P value = 7.13E6) [19], although the C-index is also slightly lower. We also detected significant survival differences in DFS using all three models, which to our knowledge has not been previously investigated. Interestingly, the subgroups from Inception model have the most significance in OS.

Another previous study performed integration of genomic data and cellular morphological features of histopathological images for clear cell renal cell carcinoma, finding that an integrated risk index from genomics and histopathological images correlated well with survival [22]. A final previous study [23] developed a CNN model using both histopathological and genomic data from brain tumors, which surpassed the current state of the art in the prediction of overall survival. Our study identified correlations between image features and biological pathways, and we expect that their integration may contribute to the better understanding of HCC etiology. Future work will involve experimenting with other CNN models, as well as further exploring the biological interpretation of our pre-trained models.

## Conclusions

The image features extracted from HCC histopathological images using pre-trained CNN models VGG16, Inception V3 and ResNet 50 can accurately distinguish normal and cancer samples. Furthermore, these image features are significantly correlated with clinical survival and biological pathways.

## Methods

### HCC datasets

We downloaded HCC histopathological images of diagnostic slides from the National Cancer Institute Genomic Data Commons Data Portal, where molecular datasets from The Cancer Genome Atlas (TCGA) Liver Hepatocellular Carcinoma such as Transcriptomics, DNA Methylation, Copy Number Variation as well as the clinical files are all available. In total, we obtained 966 HE-stained whole slide images from 421 scanned HCC subjects (421 tumor samples and 105 normal tissue adjacent to tumor samples). The images were digitized and stored in .svs files which contain pyramids of tiled images with differing levels of magnification. We used the Python modules OpenSlide and DeepZoomGenerator to read the image files. Most of the files had 3 or 4 levels of size and resolution, where level 4 indicates the largest size (median pixels: 89640 × 35870) and the highest resolution and level 3 is approximately 1/16th the size of level 4 (median pixels: 5601 × 2249.5). To reduce memory usage and processing time, we read either level 3 images or downsampled level 4 images by a factor of 16. We removed two files which were either broken or did not contain level 3 or level 4 information. In total, we used 964 files for analysis. As mentioned above, we cropped each image to either 256 × 256 or 512 × 512 pixels before extracting features.

We downloaded clinical files containing overall survival (OS) and disease-free survival (DFS) information from the cBioPortal for Cancer Genomics website (https://www.cbioportal.org/), which provides visualization, analysis and download of large-scale cancer genomics data sets. Importantly, cBioPortal includes data for the same patient cohort from which the HCC images were taken from TCGA-Liver Hepatocellular Carcinoma. When performing OS analysis, the event of interest is death (event = 1), while the censored event is being alive (event = 0). Thus, the number of days for event 1 and event 0 are the number of days until death and number of days until last contact, respectively. In DFS analysis, the event of interest is new tumor occurrence (event = 1), while the censored event is the lack of detection of a new tumor (event = 0). In this case, the number of days for event 1 and event 0 are the number of days until detection of a new tumor and number of days until last contact, respectively.

We downloaded molecular pathway information, including integrated gene expression and copy number variation data, from the Broad Institute GDAC Firehose (https://gdac.broadinstitute.org/), which provides an open access web portal for exploring analysis-ready, standardized TCGA data including the cohort from which the TCGA-Liver Hepatocellular Carcinoma image files were collected. The PAthway Representation and Analysis by Direct Inference on Graphical Models (PARADIGM) algorithm [27] predicts the activity of molecular concepts including genes, complexes and processes and measures using Integrated Pathway Ievel (IPLs) determined using a belief propagation strategy within the pathway context. Given the copy numbers and gene expression measurements of all genes, this belief propagation iteratively updates hidden states reflecting the activities of all of the genes in a pathway so as to maximize the likelihood of the observed data given the interactions in the pathway. In the end, the inferred level of a pathway reflects both the data observed for that pathway as well as the neighborhood of activity surrounding the pathway. The analysis-ready file of IPLs calculated by the PARADIGM algorithm was used for correlation analysis between image features and biological pathways.

### Image pre-processing and Feature extraction

For each of the 964 image files from 421 tumor samples and 105 normal samples, we performed stain-color normalization, adopted from previous image studies [46] [43], for both downsampled images derived from the highest resolution. From each of these images, we then generated 50 augmented images using random color augmentation. Next, we randomly selected 20 mini patches of 256 × 256 pixel and 512 × 512 pixel crop sizes from each augmented image. We fed these crops to the three pre-trained CNN models (VGG 16, InceptionV3, and ResNet 50), which generated a total of 20 sets of features. These features were then combined into one single set of features for each image. We computed the median values of extracted features across all 50 augmented images and used these as the final feature set for each sample.

Deep CNN models such as VGG 16, Inception V3 and ResNet 50, containing millions of parameters and trained for a relatively long time on large training datasets, have reached state-of-the-art performance in image recognition and classification. We used these models to extract features in an unsupervised manner to avoid the challenges of training an entire CNN model scratch. For the Inception and ResNet models, we used nodes in the second-to-last layer as features. For the VGG model, we concatenated nodes from the last 4 layers(`block2 conv2,block3 conv3, block4 conv3, block5 conv3`) as features. In each case, the CNN network weights had been pre-trained using ImageNet data [47]. We implemented the above steps using Keras, a popular Python framework for deep learning.

### Sample Visualization

To visualize samples, we first used Principal Component Analysis (PCA) the dimensionality of image features. We then applied the t-Distributed Stochastic Neighbor Embedding (t-SNE) method to visualize the first 10 components in 2 dimensions. T-SNE can reduce dimensionality of data samples based on conditional probabilities that preserve local similarity. We used a t-SNE implementation that makes Barnes-Hut approximations, allowing it to be applied on large real-world datasets [48]. We set the perplexity to 15, and sample points were colored using the group information.

### Supervised classification from image features

We applied a linear Support Vector Machine (SVM) classifier [49] to discriminate between tumor and normal samples using the image features. We used 6-fold cross validation to train the model. To evaluate classifier performance, we visualized the Receiver Operating Characteristic (ROC) curve generated using cross-validation, with false positive rate on the X axis and true positive rate on the Y axis. We calculated the Area under the ROC curve (AUC) for each cross-validation fold, as well as the overall mean value. We also plotted the 2-class precision-recall curve to visualize the tradeoff between precision and recall for different prediction thresholds. A high AUCrepresents both high recall and high precision, which translate to low false positive and false negative rates. Using average precision (AP), we summarized the mean precisions achieved at each prediction threshold. We used the Python module Scikit-learn to perform classification.

### Survival analysis

To perform univariate survival analysis for each image feature individually, we applied Cox Proportional Hazards (CoxPH) regression models using the R package ‘survival’ for both overall and disease-free survival. We used a log-rank test to select significant image features with p-value ≤ 0.05.

For multivariate survival analysis, we used the R package ‘glmnet’ to build CoxPH overall survival models based on image features from the three CNN models. We applied elastic net regularization with alpha = 0.5, which corresponds to equal parts lasso and ridge regularization. To find the best lambda, we applied 10-fold cross validation. We evaluated models with the Concordance index (C-index) and logrank test. The C-index quantifies the quality of rankings and can be interpreted as the fraction of all pairs of individuals whose predicted survival times are correctly ordered [50] [51]. A C-index of 0.5 indicates that predictions are no better than random. We fit Kaplan–Meier curves to visualize survival probabilities.

### Subgroup discovery

Using the Python module Scikit-learn, we applied K-Means clustering across all cancer samples to discover subgroups based on image features which are significant in both overall and disease-free survival. The K-Means algorithm [52] clusters samples by minimizing within-cluster sum-of-squares distances for a given number of groups, which we varied between 2-12. To evaluate the clustering results, we applied two metrics–the mean Silhouette coefficient and the Davies-Bouldin index. The Silhouette coefficient [53] takes values between −1 and 1, and it is calculated based on the mean intra-cluster distance and the mean nearest-cluster distance for each sample, where a higher value corresponds to better cluster separation. Values near 0 indicate overlapping clusters, while negative values indicate assignment of samples to the wrong cluster. The Davies-Bouldin index [54] is calculated based on the average similarity between each cluster and its most similar one, where a lower index indicates a better separation. Indices close to 0 indicate a good partition. We also constructed CoxPH models to detect survival differences between subgroups, again using C-index and log-rank test for evaluation. As before, we fit Kaplan–Meier curves to visualize the survival probabilities for each subgroup.

### Correlation between image features and pathways

We calculated the Pearson correlation between image features and pathways using the Python module scipy, which measures linear relationships between variables. Pearson correlation coefficients can range between −1 and 1, with 0 implying no correlation. Each correlation coefficient is accompanied by a p-value, which indicates the significance of the coefficient in either the positive or negative direction. To correct for multiple hypothesis testing, we adjusted p-values using the BH approach [28]. We selected significant correlations between image features and pathways as those whose adjusted p-values ≤ 0.05.

## Author’s contributions

## Acknowledgements

Text for this section …

